# Focused Ultrasound Neuromodulation: Disentangling Neural Signals from Mechanical Artifacts

**DOI:** 10.64898/2026.11.26.769554

**Authors:** Lara C. Krisst, Mohammad Shehata, Daniel Wagenaar, Shinsuke Shimojo

## Abstract

Transcranial focused ultrasound (tFUS) can noninvasively modulate neural activity in the human brain (Blackmore et al., 2019; Fomenko et al., 2018; Legon et al., 2014), and EEG, with millisecond resolution, is well suited to track the timing of cortical engagement. In humans, tFUS aimed at primary visual cortex (V1) has been reported to evoke phosphenes, with EEG peaks (N55, P100) during sonication (Lee et al., 2016). Ultrasound can also activate cortex indirectly through the auditory pathway (Sato et al., 2018), although airborne sound from the transducer can be masked (Braun et al., 2020; Liang et al., 2023). Whether the N55 and P100 peaks reflect neuronal activation or artifacts (i.e., non-neuronal signals produced by the ultrasound itself) is unclear. We set out to ask whether EEG decoding could detect neural engagement during tFUS to V1 in eight participants receiving 150 or 300-ms pulses or sham. Decoding of tFUS versus sham reached as much as 93% for exactly the duration of each pulse (cluster p = .004 and .008), with no later cluster significant. But the same pulse-locked step appeared on all channels, scaled with drive, and was also recorded from a foam head and a forehead piezo pickup, identifying it as mechanical. Decoding during sonication must therefore be interpreted as a mechanical artifact rather than cortical engagement. EEG peaks observed within a pulse cannot on their own be taken as evidence of neural engagement. EEG studies of tFUS will need controls that reproduce the vibration without focusing on the brain, such as an active sham.

## Methods

### Participants

Eight participants (*n* = 8) received monetary compensation for their participation. All participants reported no neurological disorders. The experimental protocol was approved by the California Institute of Technology Review Board, and informed consent was obtained before each session.

### Procedure

The stimuli and task are illustrated in Figure 1. All stimuli were generated in Python. Participants sat in front of a blank OLED screen at a viewing distance of 100 cm. A fixation cross was presented throughout the trial, except during the response interval. On each trial an Auditory Mondrian auditory mask, which consisted of two auditory tones were presented in succession (700; 1400ms). Transcranial focused ultrasound (tFUS) or Sham was presented at 1000 ms followed by a prompt, “Did you see any visual change?” Participants received one burst per trial, 1000 ms after trial start, and were instructed to report whether they experienced a change in visual perception. There were two active stimulation conditions (150; 300 ms duration), and Sham (no ultrasound output). Trials consisted of 150 trials (50 trials x 3 conditions) in random order. A single-element 265 kHz transducer (H-115, Sonic Concepts), coupled to the scalp through a 2-cm gel pad, delivered bursts pulsed at 1 kHz with a 50% duty cycle (0.5 ms on, 0.5 ms off) for 150 ms or 300 ms. It was aimed at right V1 (MNI +7, −76, 10), using Brainsight neuronavigation (Rogue Research) and acoustic simulation software (BabelBrain) on each participant’s MRI. For the 150-ms burst, the transducer was driven at 103.5 V peak-to-peak; for the 300ms burst at 73.9 Vpp. In free field these correspond to a peak pressure of 0.81 and 0.58 MPa, a mechanical index of 1.58 and 1.13, and a spatial-peak pulse-average intensity (Isppa) of 22.3 and 11.4 W/cm^2^. Acoustic parameters follow ITRUSST reporting recommendations (Martin et al., 2024). Both had a spatial-peak temporal-average intensity (Ispta) of about 120 mW/cm^2^. These are free-field values and do not account for the skull.

**Figure 1.**
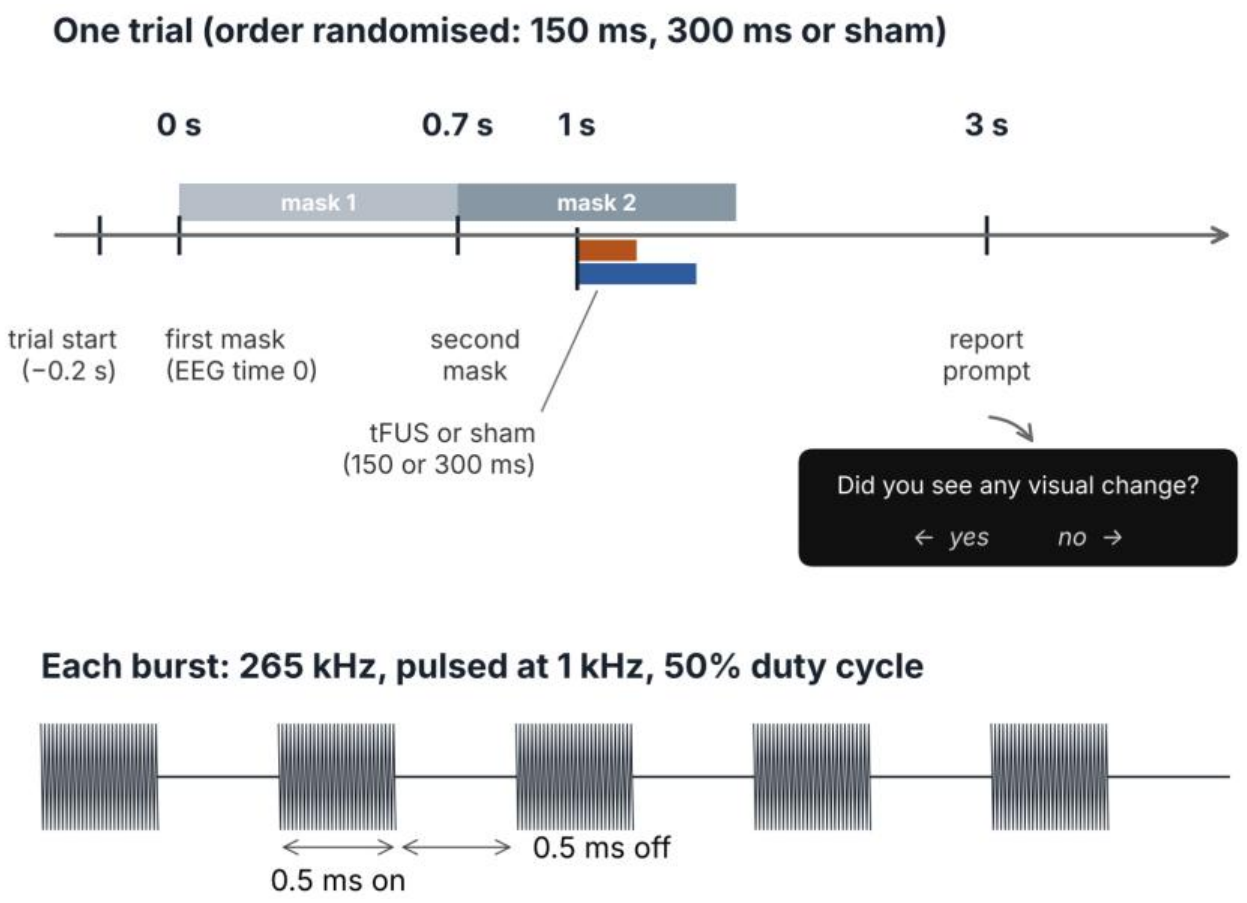
Trial structure and ultrasound burst. Each trial began with a trial-start event (−0.2 s), followed by two overlapping auditory masks starting at 0 s (EEG time 0, the first mask) and 0.7 s. A 150ms (810 kPa) or 300ms (580 kPa) tFUS burst, or sham, began at 1 s. At 3 s, participants were asked whether they had seen any visual change. Trial order was randomized. Each burst was a 265 kHz carrier pulsed at 1 kHz with a 50% duty cycle (0.5 ms on, 0.5 ms off).

### EEG Recording

EEG signals were recorded with a 128-channel BioSemi ActiveTwo system on the recording reference (CMS), with no re-referencing. Impedances were <20 kΩ. Electrodes A26–A31, B5–B8 and B12–B13 were physically removed from the cap to make room for the transducer, leaving 116 channels. A19–A25 and A32 —the strip bordering the transducer cut-out—were noisy in several participants and were also excluded. This gave a common set of 108 scalp channels for all eight participants. ICA was used only to remove blink components; stimulation-locked components were not removed. Continuous EEG was downsampled to 250 Hz, then filtered at 0.1–80 Hz, epoched from −300 to 3000 ms relative to the first audio mask, and baseline corrected from –300 ms to 0 ms from the mask. Offline analysis was performed using EEGLAB (Delorme & Makeig, 2004) and ERPLAB (Lopez-Calderon & Luck, 2014). The epochs for each condition were then averaged together.

### Decoding Analysis

Decoding used a four-dimensional matrix for each participant: time points (825) × conditions (2) × trials (~50 per condition) × electrode sites (108). Support vector machines (SVMs) with error-correcting output codes (ECOC; Dietterich & Bakiri, 1994) classified active versus sham from the distribution of the ERP signal across the 108 scalp electrodes. Decoding was performed independently at every time point (4 ms spacing) from −300 to 3000 ms. We used three-fold cross-validation, iterated 100 times: trials in each condition were randomly divided into three groups of trials (3 groups × 2 conditions × 108 electrodes at each time point), two groups serving as the training set and one as the test set. To improve signal-to-noise ratio, trials within each group were averaged, yielding three ERP waveforms per condition, and decoding was applied to these averages rather than to single trials (Bae & Luck, 2018). A separate classifier (two SVMs) was trained for each condition, and their outputs were combined into a single decision about the most likely condition for each test average. A test case was scored correct only if the classifier chose the correct condition of the two (chance = 0.5). This was repeated three times at each time point, with each group serving once as the test set, and the whole procedure was iterated 100 times with trials re-shuffled into groups before each iteration. Accuracy was averaged across the two classes, cross-validation folds and iterations to give a percentage decoding accuracy at each time point. The 150-ms and 300-ms conditions were each decoded separately against sham.

### Statistical Analysis of Decoding Accuracy

Group-level decoding accuracy was tested against chance (0.5) separately for the 150-ms and 300-ms conditions, using a cluster-based sign-flip permutation test across participants (n = 8) over 0–3000ms from the first audio mask; the pre-stimulus baseline was not included in the test. At each time point we computed a one-sample t statistic on participants’ accuracy minus 0.5 (df = 7). Adjacent time points exceeding a one-sided cluster-forming threshold (p < .05) were grouped into clusters, and each cluster’s mass was the sum of its t values. The null distribution of the maximum cluster mass was built from all 256 (2^8^) possible sign flips of participants’ accuracy-minus-chance values, and a cluster was significant if its mass was exceeded by fewer than 5% of the null values (cluster-level p < .05, one-sided). This controls the familywise error rate across time points.

## Results

### Decoding tracked the pulse

Before stimulation, decoding of tFUS versus sham was at chance (mean accuracy 0.48–0.49). Accuracy rose sharply at pulse onset and stayed elevated for the duration of the pulse, peaking at 93% for the 150-ms pulse (Fig. 2). It returned to chance within tens of milliseconds of pulse offset. The cluster-based permutation test found one significant cluster for each pulse length. For the 150 ms pulse it ran from 1004 to 1172 ms (p = .004), and for the 300-ms pulse from 1012 to 1312 ms (p = .008). Each cluster began at pulse onset (1000 ms) and ended at or just after pulse offset (1150 and 1300 ms). Outside these windows no cluster reached significance in either condition.

**Figure 2.**
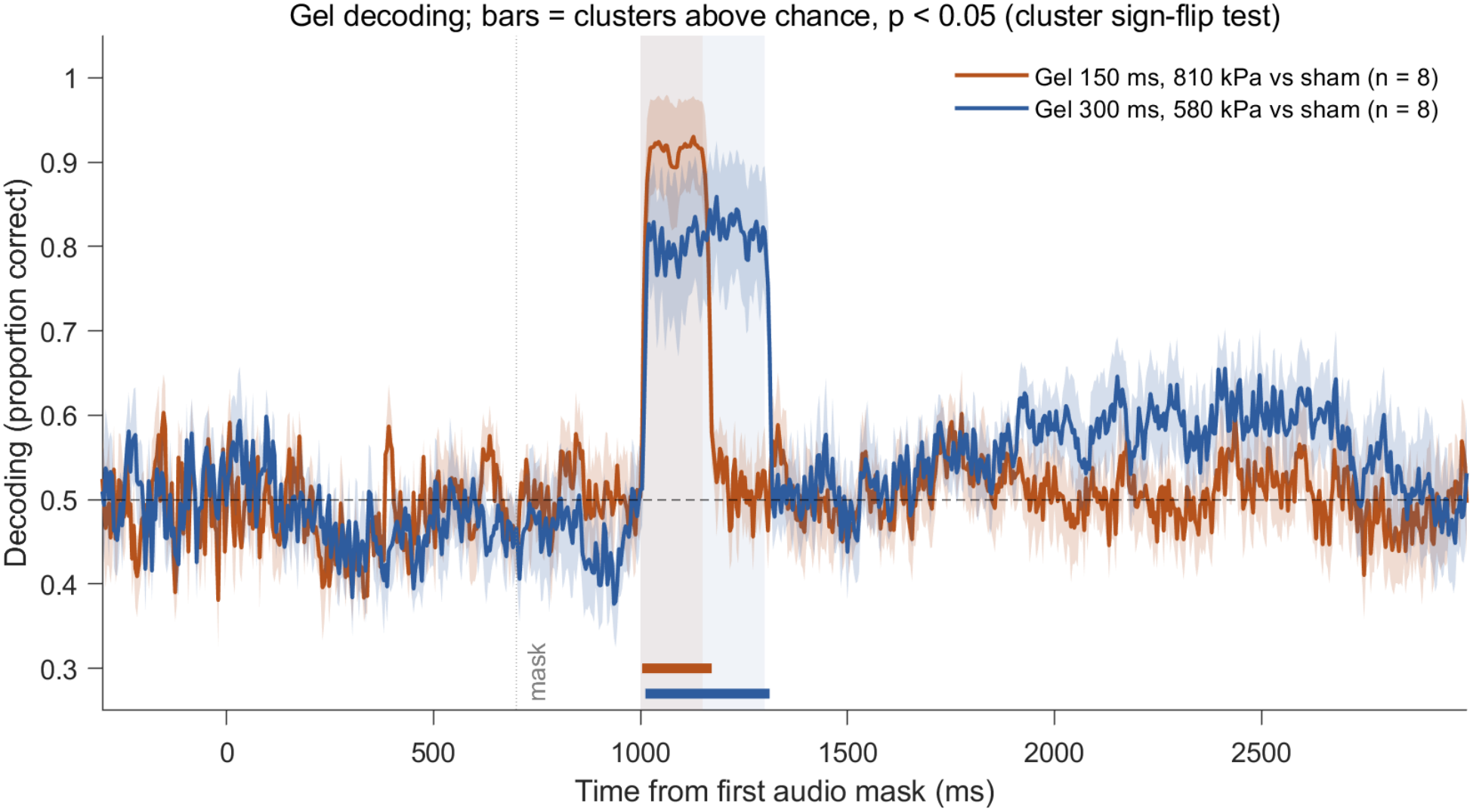
Time-resolved decoding accuracy (mean ± SEM across participants, n = 8) for the 150-ms (orange) and 300-ms (blue) conditions, each decoded against sham. Time is from the first audio mask (0 ms); tFUS or sham began at 1000 ms. Shaded bands mark the pulse windows. The dashed line is chance (0.5). Horizontal bars mark the significant clusters (cluster-based sign-flip permutation test against chance over 0–3000 ms, cluster-level p < .05, one-sided): 1004–1172 ms (150 ms, p = .004) and 1012–1312 ms (300 ms, p = .008). No other cluster was significant.

**Figure 3.**
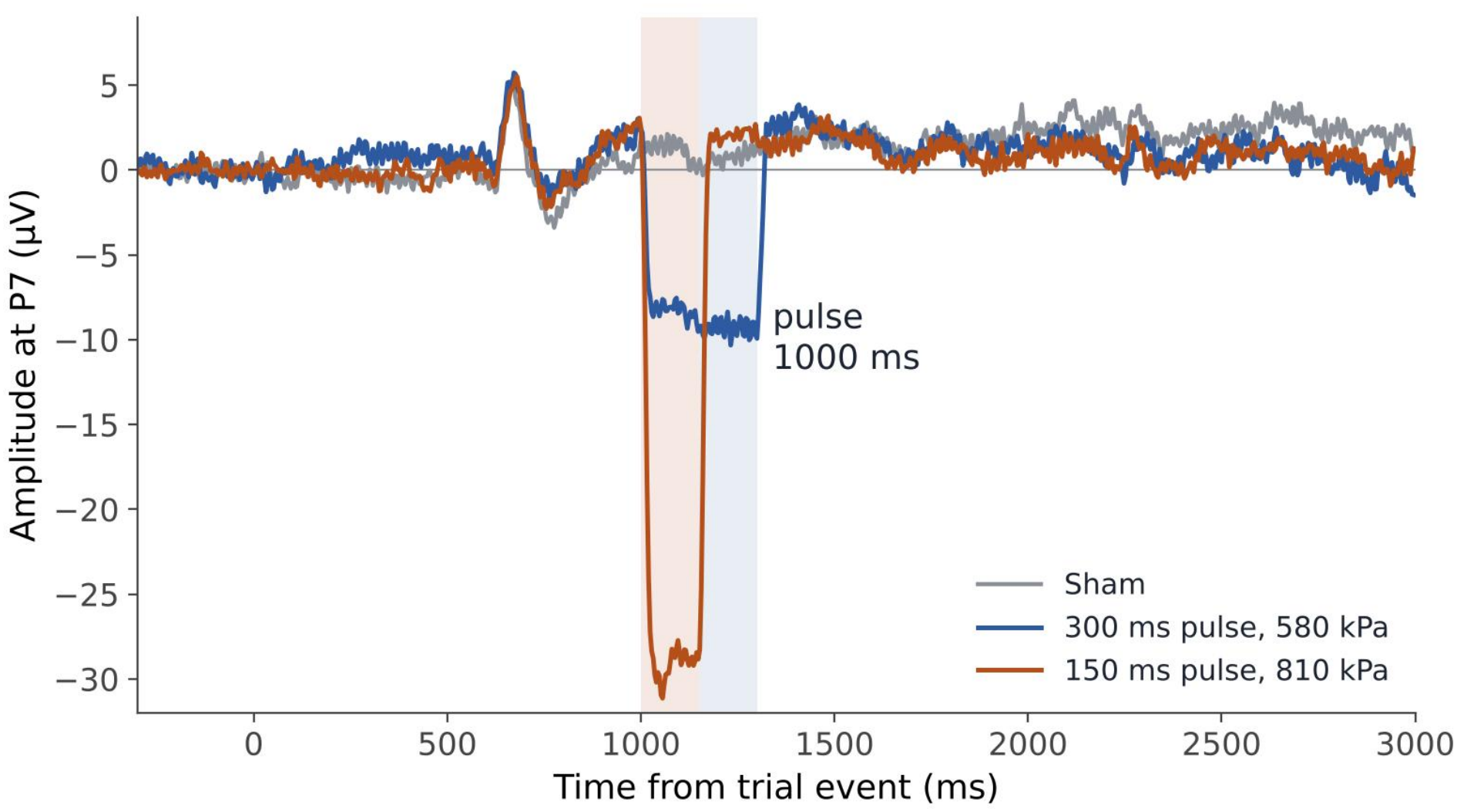
Grand-average ERP (n = 8) at electrode P7 for the 150-ms pulse (orange), the 300-ms pulse (blue) and sham (grey). Time relative to the start of first audio mask (0 ms). Shaded bands mark the 150-ms and 300-ms pulse windows beginning at 1000 ms. Both active conditions show a box-shaped negative step that begins and ends with the pulse. It measures about −30 µV for the 150-ms pulse and about −10 µV for the 300-ms pulse, and it is absent on sham. All three conditions share the small deflection around 650–700ms that follows the auditory masks.

### The pulse-locked signal is mechanical

The ERP showed a box-shaped step that began and ended with the pulse (150 or 300 ms) and was absent on sham. At P7 it measured ≈ −30 µV for the 150-ms pulse and ≈ −10 µV for the 300-ms pulse. It was negative on all 108 channels, with a similar size over the left and right hemispheres (median across channels ≈ 28 and 10 µV), so P7, which lies in the left hemisphere opposite the target, was a typical channel rather than a special one. The same step appeared when a cap was placed on a foam head phantom, and a piezo pickup on the forehead recorded the pulse.

## Discussion

We found that decoding during the pulse that could mistakenly have been interpreted as cortical engagement, instead reflected an artifact. The pulse-locked step was present on every channel, absent on sham, scaled with drive, and appeared on a foam head and a forehead piezo pickup.

The step is not an auditory response, because it also appears on a foam head, which has no auditory system. Several limitations apply: the sample is small (n = 8), only one hemisphere was targeted. How the signal reaches the electrodes is also unresolved. Next steps are to add an active sham that reproduces the vibration and sound without focusing on the brain. A time-locked, dose-dependent signal of this kind can be mistaken for a neural response, so foam-head controls should be a required part of any study that reports EEG peaks during sonication.

